# Systematic Profiling of Ale Yeast Protein Dynamics across Fermentation and Repitching

**DOI:** 10.1101/2023.09.21.558736

**Authors:** Riddhiman K. Garge, Renee C. Geck, Joseph O. Armstrong, Barbara Dunn, Daniel R. Boutz, Anna Battenhouse, Mario Leutert, Vy Dang, Pengyao Jiang, Dusan Kwiatkowski, Thorin Peiser, Hoyt McElroy, Edward M. Marcotte, Maitreya J. Dunham

## Abstract

Studying the genetic and molecular characteristics of brewing yeast strains is crucial for understanding their domestication history and adaptations accumulated over time in fermentation environments, and for guiding optimizations to the brewing process itself. *Saccharomyces cerevisiae* (brewing yeast) is amongst the most profiled organisms on the planet, yet the temporal molecular changes that underlie industrial fermentation and beer brewing remain understudied. Here, we characterized the genomic makeup of a *Saccharomyces cerevisiae* ale yeast widely used in the production of Hefeweizen beers, and applied shotgun mass spectrometry to systematically measure the proteomic changes throughout two fermentation cycles which were separated by 14 rounds of serial repitching. The resulting brewing yeast proteomics resource includes 64,740 protein abundance measurements. We found that this strain possesses typical genetic characteristics of *Saccharomyces cerevisiae* ale strains and displayed progressive shifts in molecular processes during fermentation based on protein abundance changes. We observed protein abundance differences between early fermentation batches compared to those separated by 14 rounds of serial repitching. The observed abundance differences occurred mainly in proteins involved in the metabolism of ergosterol and isobutyraldehyde. Our systematic profiling serves as a starting point for deeper characterization of how the yeast proteome changes during commercial fermentations and additionally serves as a resource to guide fermentation protocols, strain handling, and engineering practices in commercial brewing and fermentation environments. Finally, we created a web interface (https://brewing-yeast-proteomics.ccbb.utexas.edu/) to serve as a valuable resource for yeast geneticists, brewers, and biochemists to provide insights into the global trends underlying commercial beer production.

## Introduction

*Saccharomyces cerevisiae,* or brewing yeast, is used in a wide range of commercial processes including beverage fermentation, baking, biofuel generation, and pharmaceutical manufacture. The ease of culture and genetic manipulation of *S. cerevisiae* has also made it one of the most profiled organisms in academic research, including being the first eukaryote to have its genome sequenced^1^. Brewing yeast has served as a model organism to understand fundamental cellular and molecular processes and continues to provide valuable insights into human health and disease^2–7^. Despite its prevalence in laboratory settings, the systematic profiling of yeast in commercial contexts is less common.

These commercial contexts are relevant not just for understanding more about the natural history of yeast, but also due to their economic and cultural importance. The global beer market was worth $744 billion in 2020 and is expected to grow to $768 billion in 2023^8^. While research on brewing yeasts has been performed over many decades, new techniques – especially in genomics, proteomics, and related high throughput profiling – allow us to better understand the genetic and molecular changes that have occurred in yeast during brewery fermentations.

Throughout brewing and fermentation, yeast must cope with diverse and changing stressors, including fluctuations in nutrient and ethanol levels, oxygen availability, and temperature^9^. At the start of the brewing process, yeast is inoculated into a fermentation vessel containing aerated wort, a cooled aqueous extract containing the sugars from boiled malted grains along with the aromatic and bittering compounds from hops. Once the yeast cells adapt to the new environment during the lag phase, they begin to grow exponentially and rapidly deplete the available oxygen creating an anaerobic environment. Along with oxygen, sugars and other essential nutrients are depleted, with concomitant production of ethanol, all of which stress the yeast over the course of fermentation. As the yeast adapt to these successive stressors during the brewing process, cells rapidly shift their gene expression profiles, leading to changes in protein and metabolite levels that help them survive^9^.

High-throughput approaches for parallel cell-wide measurement of different classes of cellular molecules, such as DNA sequencing (genome), mRNA sequencing (transcriptome), protein profiling (proteome) and metabolite profiling by mass spectrometry (metabolome), offer a path to deeper understanding of how yeast cells respond to the brewing environment. Hundreds of brewing strains, including both *S. cerevisiae* ale yeasts and *S. pastorianus* lager yeasts (which are interspecific hybrids between *S. cerevisiae* and *S. eubayanus*) have had their full genomes sequenced, yielding insights into domestication history and genetic differences related to flavor and style^10–12^. Gene expression profiles during growth in wort have been characterized for ale yeasts and lager yeasts in commercial brews and in small wort fermentations^13, 14^. Investigators have also characterized ale yeasts’ proteomes^15^ and metabolomes^13, 16^ during wort fermentation, although high-throughput proteomics based studies are infrequent. The majority of these previous brewing yeast multi-omics studies focused on the analysis of the proteins and metabolites known to be involved in the production of flavor components such as esters and higher alcohols by sampling beer or wort. However, few studies have sampled the actual brewing yeast populations to understand the global aspects of how yeast proteomes temporally change over time during commercial fermentation.

One common commercial brewing condition for which there have been few comprehensive ale yeast studies is “serial repitching”, a process in which brewers collect yeast cells at the end of a fermentation cycle and use it to inoculate (or “pitch”) a new batch. Serial repitching is mainly done for ease and efficiency of brewing and preserving the sensory or taste profile of the beer. The number of repitches varies across breweries, type of fermentation, and strain of yeast used. However, excessive rounds of repitching can adversely affect yeast fermentation and the taste profiles of the final product^17^. Serial repitching results in repeated exposures to physical, biological, and chemical stresses, which can lead to both reversible and irreversible damage to the yeast cells, with progressive loss in cell viability occurring with increasing pitch number^17^. Plasma membrane damage and stress-activated gene expression programs that result in increased glycogen accumulation and intracellular trehalose levels increase with each subsequent repitch^18^. Beer quality can be adversely affected due to changes in flavor and aroma profiles, decreased viability of yeast cells, and the increased likelihood of undesired microbial contamination, and are among the major reasons that brewers stop repitching after about 8-10 batches and restart brewing with a fresh yeast culture.

Genetic changes in the yeast during repitching can contribute to the altered profiles and viability. Previous studies have characterized mutations and traits that accumulate in ale yeast over repitching^9, 18–21^. For instance, one study observed repeated changes in copy number of chromosome V and mitotic recombinations that changed allele balance and subtelomeric gene copy number at regions on chromosomes VIII and XV^19^. Changes in yeast traits such as flocculation were also observed, though not linked definitively to these mutations. However, changes in gene expression, protein levels, and metabolite abundance were not measured.

While there have been many ecological studies tracing the origin, evolution, and physiology of fermentation and brewing strains using genomics^10–12^, few have focused on molecular changes at the protein level and across brewing cycles^22–25^. To gain a more comprehensive understanding of the molecular changes associated with brewing, we characterized the genetic features of a Hefeweizen ale yeast and measured temporal proteomic changes across two fermentation cycles separated by serial repitching. Across 64,740 protein abundance measurements, we found many processes altered over fermentation in both time courses: in particular, we observed drastic changes in yeast proteomes across the first two days of fermentation largely dominated by ribosome biogenesis and translation. Additionally, we cataloged unique changes between the two fermentation batches, observing that lipid and sterol biosynthetic processes were upregulated in the later batch. This dataset serves as a foundational resource to finely characterize the molecular changes underlying commercial ale fermentation and offers a starting point to perturb, modify, or engineer flavor and strain characteristics in commercial and craft brewing settings.

## Results

### Hefeweizen yeast shares genetic characteristics with other ale brewing strains

In this study, we used Wyeast 3068, a Weihenstephan Weizen tetraploid *S. cerevisiae* ale strain^26^ frequently used to brew Hefeweizen style beers. This popular German style wheat beer strain^27^ has previously been whole genome sequenced and ecologically annotated^11, 12^, allowing for comparisons against existing resources. Moreover, the relatively short fermentation cycles in Hefeweizen brewing (4-6 days) allowed us to sample yeast populations across brewing cycles separated by 14 cycles of serial repitching.

We confirmed the identity and analyzed the genetic characteristics of our strain using whole genome sequencing and alignment to the S288C *Saccharomyces cerevisiae* genome. Our data demonstrated mostly even coverage across the yeast genome (Sup. Fig. 1A), with depth differences corresponding to copy number changes (Sup. Fig. 1B). Several genes found in the reference laboratory strain were deleted in Wyeast 3068 (Sup. Table 1). Deleted genes were enriched for processes of flocculation (p≤4.59×10^-5^, *FLO5, FLO9, FLO1, FLO10*), carbohydrate transmembrane transport (p≤2.99×10^-3^, *HXT15, HXT16, MPH2, MPH3, AQY3*), asparagine catabolism (p≤3.34×10^-6^, *ASP3-3, ASP3-2, ASP3-1, ASP3-4*), and transposition (p≤2.30×10^-4^, *YIL082W-A, YPL060C-A, YLR157C-A, YGR109W-A, YGR109W-B, YJL113W, YJL114W, YLR157C-B*), and some of these genes have previously been found to be deleted in ale strains^11, 12^. Also like other ale strains, Wyeast 3068 was largely tetraploid with several chromosomal copy gains and losses, most notably possessing an extra copy of chromosome V and losing a copy of chromosome X (Sup. Fig. 1B). Additionally, a majority of the chromosomes contained genomic regions that had not changed in copy number, but had experienced a loss of heterozygosity (Sup. Fig. 1C), as seen in other ale strains^28^.

### A global view of temporal protein changes across brewing

To understand protein dynamics during brewing cycles, we periodically sampled yeast populations directly from the fermentation tank over a brewing time course inoculated with a freshly prepared stock of yeast (hereby termed Batch 1). To study the effects of repitching on yeast proteomes, we similarly sampled yeast after 14 repitches (Batch 15), with each pitch (fermentation) using a similar Hefeweizen wort composition. As typical for brewery repitching, the yeast cells from the previously finished fermentation were directly inoculated into the subsequent fermentation without any outgrowth. For Batch 1 and Batch 15, we collected samples representing near matched timepoints across the four days of brewing. To achieve this, we collected beer and isolated yeast populations directly from the fermentation tank. We also collected samples 24 hours post-crash (PC), when the fermentations were chilled or “cold-crashed” to sediment the yeast (Fig. 1). After the cold crash, the majority of the yeast cells were separated from the beer. The beer was moved to a conditioning tank to proceed with maturation where some additional yeast was sedimented (not to be harvested) and the beer flavor matured and turbidity was homogenized. The majority of yeast cells that were harvested from the cold crash were held at 4.4°C for 1-2 days until they were used to inoculate the next batch. For Batch 15, we additionally sampled the residual yeast in the conditioning tank over three days after the Batch 15 fermentation cycle, hypothesizing that the proteomes of yeast in the conditioning tank would represent different physiological states to those in the fermentation tank. In total, we sampled 17 time points: seven from Batch 1 and ten from Batch 15 (Sup. Table 2). Independent duplicate samples were collected and processed from each timepoint.

**Figure 1.**
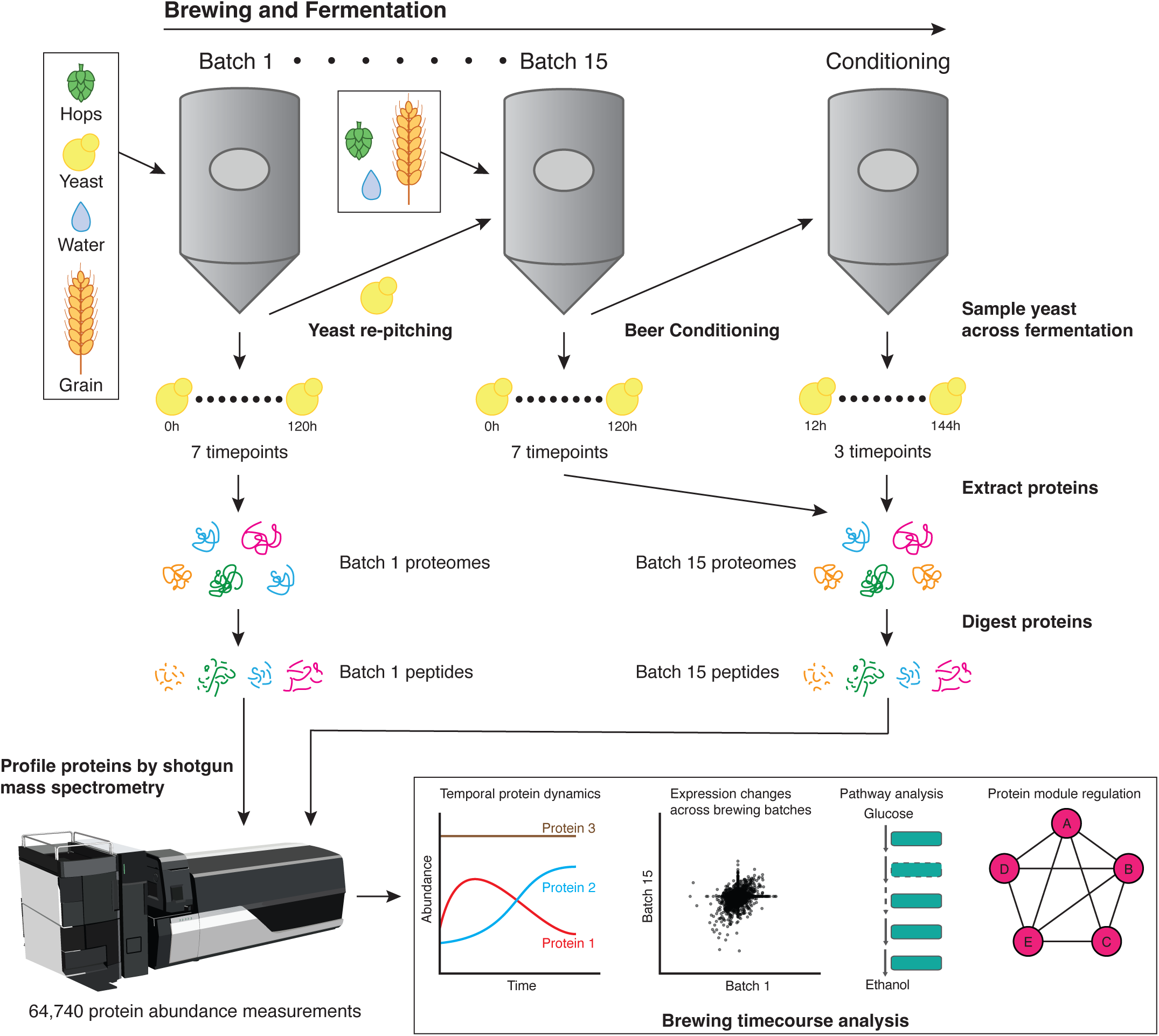
Overview of Wyeast 3068 fermentation analysis via mass spectrometry. To systematically profile the proteomic changes across fermentation, we sampled 2 commercial production scale fermentation batches (Batches 1 and 15) consisting of the same parental strain population. Batches 1 and 15 consisted of 7 time points each, with each successive time point being no more than a few days apart. At the end of fermentation in Batch 15, beer fermentations were subjected to conditioning during which we sampled 3 additional time points. To compare protein dynamics between Batch 1 and 15, 6 time points were correspondingly matched to investigate batch-specific protein changes.

We then performed high-throughput shotgun mass spectrometry on the yeast sampled from each batch (see Methods) to generate proteomic snapshots of all assayed yeast proteins and their abundance over rounds of brewing. For every protein, we computed three abundance metrics: parts per million (ppm)^29^, intensity-based absolute quantification (iBAQ), and label-free quantification (LFQ) values^30–32^ (see Methods). Since the LFQ values had the highest correlation between replicates and were corrected for technical variation between samples, we proceeded with LFQ abundances for all subsequent analyses.

In total, we made 64,740 protein abundance measurements for over 2600 proteins detected in at least one time point, out of a total of 5610 possible experimentally detectable yeast proteins^33^. We compared the correlation between replicates for each time point by plotting the LFQ intensity values for every protein detected in that time point and found that replicates of a particular time point in one batch were highly correlated, with Pearson correlation coefficients ranging from 0.84 to 0.94 (Sup. Fig. 2A). We generated a similarity matrix by performing all-by-all Pearson correlation calculations across the 34 samples and observed that replicates of a single time point within a particular batch were more correlated to each other than to any other time point (Sup. Fig. 2B). We observed three groups with high correlation: two composed of time points within Batch 1 and Batch 15, respectively, and a third group of matched time points across both batches. Despite high correlation, we found that the replicates of the earliest samples, 6h from Batch 1 and 3h from Batch 15, were the least correlated with other time points assayed, suggesting that the proteome profiles of the earliest fermentation time points were most different compared to the rest of the fermentation.

Since DNA copy number differences can alter protein levels^34^, we investigated whether the observed copy number alterations affected initial protein levels. As expected, the protein products of genes that were deleted (Sup. Table 1) were not detected at the protein level. Among detected proteins, there was no correlation between read coverage and protein abundance at the initiation of Batch 1 on a per-gene/protein basis (Sup. Fig. 1D, Sup. Table 3, adjusted R^2^=0.00008246, p≤0.2703), nor by comparing the average read depth per gene to the average abundance of the corresponding protein product for all genes on each chromosome (Sup. Fig. 1E, adjusted R^2^=- 0.04194, p≤0.5392). There was also no correlation at any of the later time points (Sup. Tables 3-4). Given this lack of correlation, we did not normalize the mass spectrometry data to the gene copy number inferred from sequencing read depth.

Given the high correlation between replicates, we summed the intensity values across both replicates to maximize the number of proteins detected in the dataset (Sup. Fig. 2A-B, see Methods). On average, 368 proteins were detected in one replicate but not the other (Sup. Fig. 2C, Sup. Table 5). We next summarized our dataset to collapse protein measurements by batches. In total, we detected 2518 proteins in Batch 1, 2504 proteins in Batch 15, and 2280 proteins in the conditioning tank (Sup. Fig. 2D). We found that 2274 proteins were common to both Batches 1 and 15, while the conditioning tank sampled 231 fewer proteins. We observed 44 proteins exclusive to Batch 1, 16 proteins exclusive to Batch 15, and 6 proteins specific to the conditioning tank.

To uncover the trends underlying fermentation, we normalized LFQ intensity values for every protein to the mean abundance across all sampled time points (Sup. Table 6) and performed hierarchical clustering and GO analysis on all proteins that changed at least two-fold. We identified clusters by node depth (Sup. Table 7, see Methods) for which we annotated the biological processes, molecular functions, and cellular components by gene ontology analysis using ClusterProfiler^35^ (Fig. 2, Sup. Table 8). We found generally consistent expression trends between batches, with protein abundances similarly changing across time in both batches. Specifically, both batches have an initial increase in ribosome biogenesis that decreases after the first day of fermentation (Fig. 2, cluster 1b), whereas proteins involved in carbohydrate and lipid metabolism and oxidation tended to have low abundance early in fermentation and increased over time (Fig. 2, cluster 7b). These changes were expected as the yeast cells are rapidly dividing over the first two days of fermentation, exhausting the preferred carbon sources. The protein levels in the Batch 15 post-fermentation conditioning tank samples were similar to the later fermentation time points in Batch 15, with the exception of enrichment for mitochondrial gene expression (Fig. 2, cluster 7c) and another cluster with proteins strongly downregulated in the final conditioning time point (Fig. 2, cluster 3), which was not significantly enriched for GO terms.

**Figure 2.**
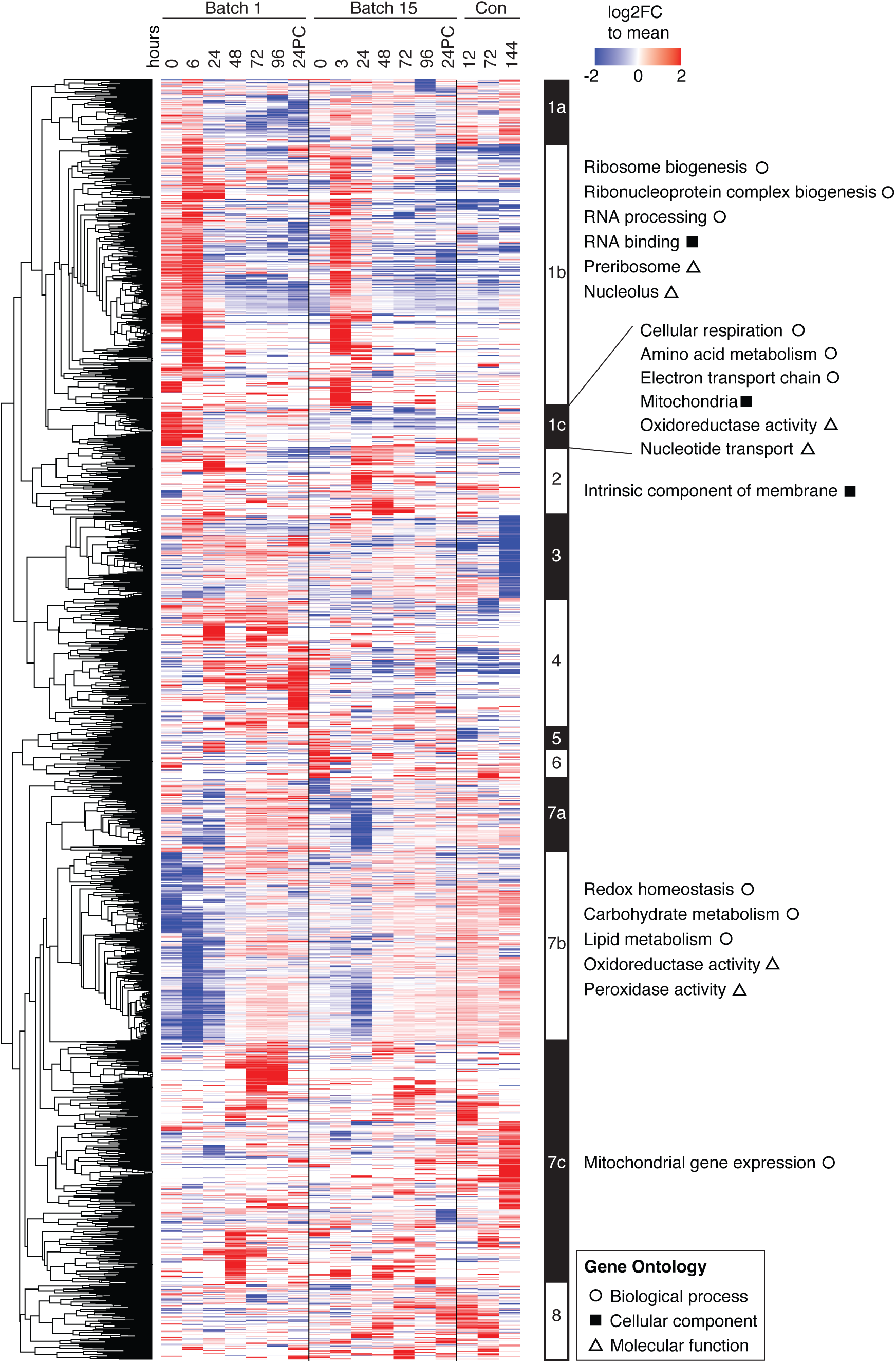
A global view of ale yeast protein changes across brewing cycles. Heatmap depicting changes in abundances of 1891 proteins which displayed at least a two-fold change relative to the mean across all timepoints (rows) over both brewing batches and final conditioning (columns). Values are colored as log_2_ fold change relative to the row mean normalized abundance (see Methods). Proteins contained within clusters (defined by node depth; see Methods) were annotated from ClusterProfiler^35^ filtering terms to an adjusted p-value (Benjamini-Hochberg correction) of <0.05 and a multiple hypothesis testing FDR of 5%. Each cell is the sum of two replicates with column time points indicated in hours. 24PC indicates 24 hours post cold crashing after fermentation.

### Brewing yeast proteomes change drastically in the first 24 hours of brewing

Yeast strains undergo complex molecular changes while they adapt to both physical and chemical changes including nutrient deprivation, loss of oxygen, and increased alcohol production over time. Previous studies have profiled the metabolites, gene transcripts, and proteins present during beer fermentation, but very few studies have performed analyses comparing different times across a complete fermentation cycle^15, 23, 25, 36–38^. We sought to understand how yeast protein expression patterns change between any two time points in the fermentation tank. Since LFQ abundances are log-normally distributed, we log_2_ transformed the LFQ abundance values in each time point, subtracted the median abundance across the time point (to equalize variances), and performed differential expression testing across time point pairs. Comparing time points all-by-all, we found that the number of differentially expressed proteins (DEPs) ranged from two to 618, with an average of 195 between any two time points (Fig. 3A, Sup. Table 9). When comparing consecutive time points, we observed the largest number of DEPs between early time point pairs in both Batch 1 and Batch 15, which progressively decreased over time (Fig. 3B, Sup. Table 9).

**Figure 3.**
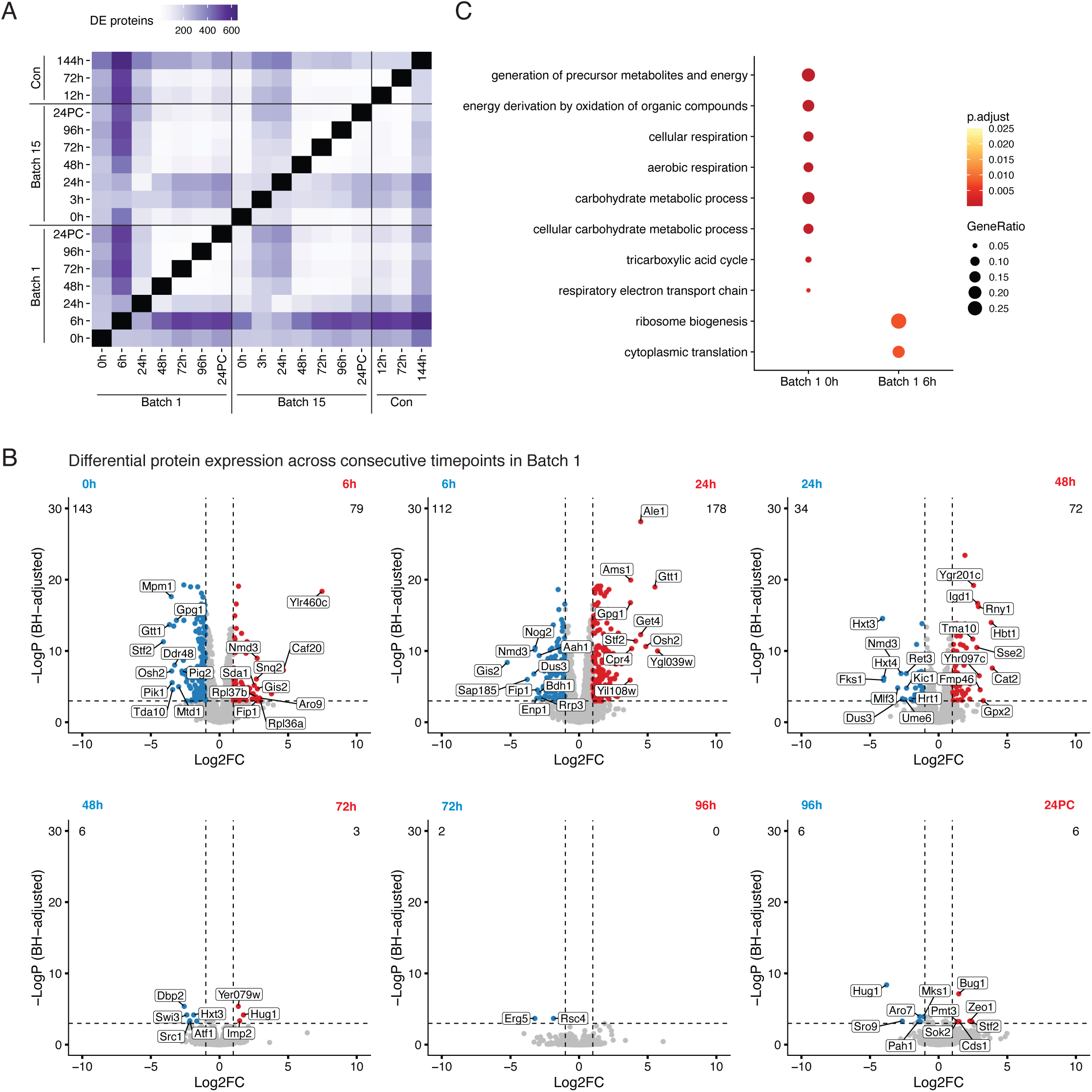
Differential protein expression dynamics across brewing cycles. A) Heatmap representing the number of differentially expressed proteins (DEPs) across pairs of fermentation time points, indicated in hours; 24PC indicates 24 hours post cold crashing after fermentation. DEPs were determined by a standard t-test. B) Volcano plots determining the protein expression changes across consecutive Batch 1 time points. C) Dot plot representing the top enriched Gene Ontology terms between the 0h and 6h time point in Batch 1.

To elucidate the processes that are temporally regulated throughout fermentation, we performed GO analysis using the DEPs detected between time points. When comparing the first 2 consecutive timepoints (Batch 1 0h and 6h) we detected 222 DEPs (Sup. Table 9) and observed an enrichment at the beginning of Batch 1 for proteins involved in aerobic respiration, generation of precursor metabolites and energy, and cellular response to stress. On the other hand, proteins detected after six hours of fermentation were significantly enriched for processes related to ribosome biogenesis and translation (Fig. 3C, Sup. Table 10). After the first day of brewing, the proteins involved in these processes along with rRNA regulatory processes were downregulated (Sup. Table 10).

We also wanted to compare how protein levels differed between batches. Protein levels in matched time points between Batch 1 and 15 were generally well correlated (Sup. Fig. 3A). We compared the initial time points of both batches and found 213 DEPs (Sup. Fig. 3B). Aerobic respiration and amino acid biosynthesis processes were enriched in Batch 1 DEPs, and sterol and lipid biosynthesis processes were enriched in Batch 15 DEPs (Sup. Fig. 3C, Sup. Table 10). Given the high correlation between protein levels in the early Batch 1 (6h) and Batch 15 (3h) time points (Sup. Fig. 3D), we also compared processes between these time points (Sup. Fig. 3D) and observed that Batch 15 was enriched for sterol biosynthesis and glycogen and cellular alcohol metabolic processes (Sup. Table 10). Though we observed 116 DEPs between the batches 24 hours post crash (Sup. Fig. 3B), no significant GO terms were enriched for this set of differing proteins between the batches at this time point.

### Uncovering the temporal changes in central metabolic proteins across brewing

While we observed global temporal protein changes over fermentation using GO analysis, we wanted to better understand the degree to which these changes were coordinated by investigating biochemical pathways. We first focused on the two main pathways involved in yeast fermentation – glycolysis and the Tricarboxylic Acid (TCA) cycle – as their regulation is central to alcohol production (Sup. Fig. 4A). On calculating the pairwise Pearson correlation of LFQ values across all samples for all pairs of proteins in the glycolysis and TCA cycle pathways, we found that sets of proteins in a given pathway tended to be well correlated. Generally, proteins involved in the TCA cycle were more correlated to each other than those involved in glycolysis (Sup. Fig. 4A). Intriguingly, however, we also found many instances of correlation between the proteins involved in glycolysis and those involved in the TCA cycle. Of the 23 glycolytic proteins detected, Hxk1, Tdh1, Adh4, Pgk1, Gpm1, Eno2, Eno1, and Cdc19 were highly correlated with enzymes in the TCA cycle (Sup. Fig. 4A).

When considering changes across fermentation, we expected glycolytic proteins to have high abundance early in the fermentation cycle and gradually decrease as sugar was consumed. However, we found that trends in the glycolytic proteins were often noisy (Fig. 4A). The levels of proteins involved in the TCA cycle were in line with the expected profile of a strain in a low-sugar and anaerobic environment^39^: TCA cycle enzymes were downregulated over fermentation cycles across both batches and only increased their abundances during the conditioning phase, when yeast were exposed to additional oxygen while being moved to a conditioning tank (Fig. 4B).

**Figure 4.**
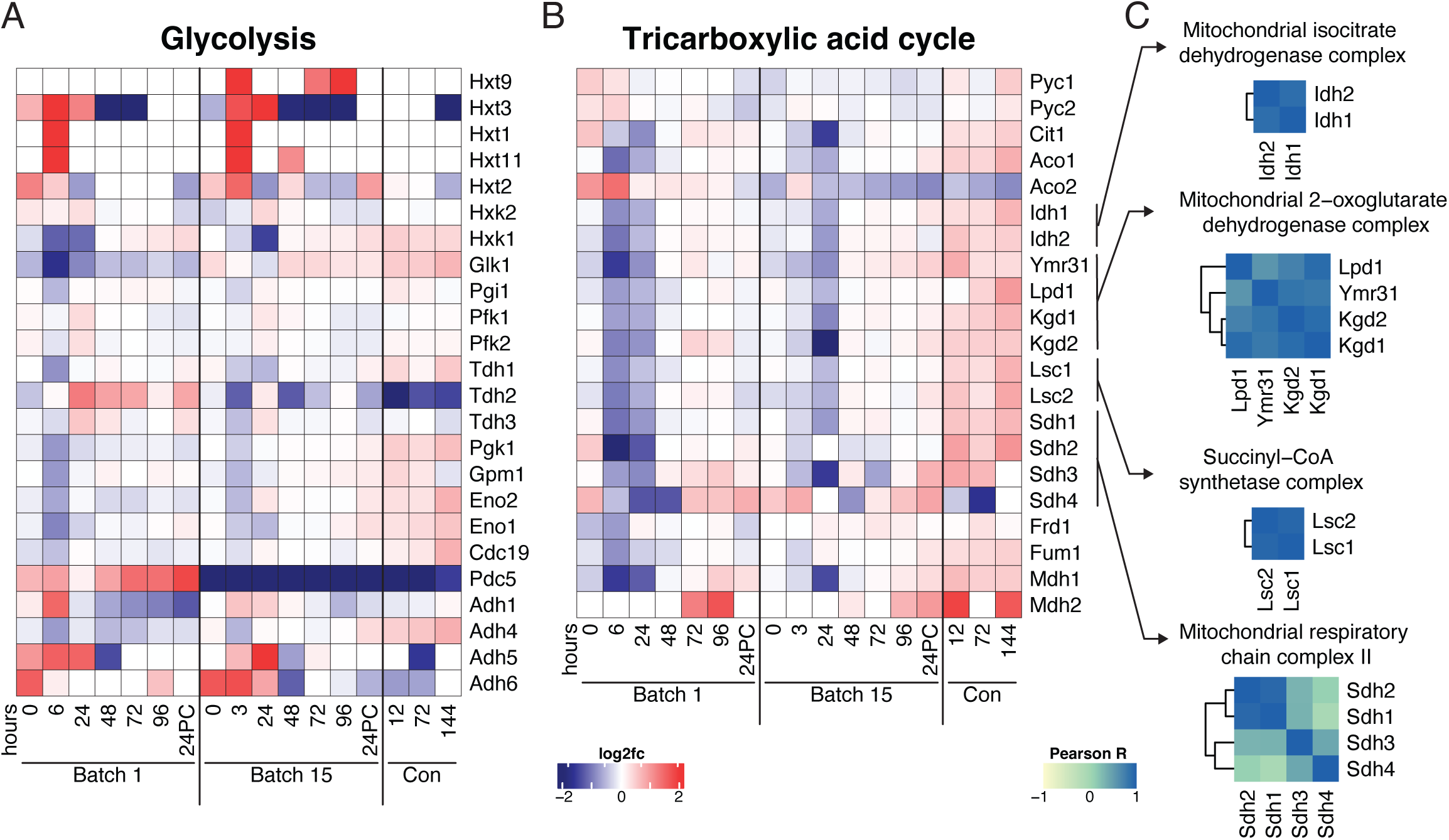
Systems-level regulation within protein modules across brewing. Heatmaps depicting the expression profiles of proteins (rows) across fermentation time (columns) in A) Glycolysis and B) the Tricarboxylic Acid (TCA) Cycle. Proteins in heatmaps ordered according to biochemical steps in their respective pathways C) Pairwise Pearson correlations across proteins within complexes involved in the TCA cycle.

We observed similarities between batches for the abundance profiles over time of enzymes catalyzing subsequent steps in the TCA cycle, demonstrated by the high degree of correlation between members of the pathway (Sup. Fig. 4B). Since enzymes in the TCA cycle interface together to form complexes, we reasoned that enzyme complexes might be coregulated throughout fermentation. Generally, we found this to be the case, with high correlation among members of the isocitrate dehydrogenase, 2-oxoglutarate dehydrogenase, and succinyl-CoA synthetase complexes (Fig. 4C).

Generally, protein profiles in glycolysis and the TCA cycle were similar between batches (Fig. 4A-B). One striking exception to this was a minor isoform of pyruvate decarboxylase, Pdc5, which converts pyruvate to acetaldehyde. Pdc5 was elevated in Batch 1 compared to its mean abundance across the dataset while, in Batch 15 we saw the opposite trend. (Fig. 4A, Sup. Fig. 4B). Although peptides unique to major isoform Pdc1 were not detected, we detected Pdc1 as a part of a protein group along with Thi3. The abundance of this group did not differ between batches or along the time course (Sup. Fig. 4B). Other proteins that differed between batches included pyruvate carboxylases Pyc1 and Pyc2 and aconitate hydrolases Aco1 and Aco2. Pyc1 and Pyc2 had higher levels for the first six hours of fermentation in Batch 1, but displayed the opposite trend in Batch 15, indicative of strain adaptation to fermentation conditions. Despite acting on the same substrate, Aco1 and Aco2 exhibited unique profiles: Aco1 levels increased across each batch whereas Aco2 decreased, and was higher in Batch 1 than Batch 15.

We also examined proteins in other pathways related to pyruvate metabolism (Sup. Fig. 4C). Pyruvate dehydrogenases Pda1 and Pdb1, which shunt carbons to the TCA cycle, maintained stable levels across fermentation time and between batches (Sup. Fig. 4B). Alcohol dehydrogenases exhibited varying profiles: the abundances of Adh1 and Adh4 remained relatively unchanged between batches and across fermentation, while Adh5 and Adh6 had lower abundances that further decreased after one to two days of fermentation (Sup. Fig. 4B). Bdh1, which converts 2,3-butanediol to acetoin, was markedly lower in abundance in Batch 15 and decreased over time.

To take an unbiased approach to identify the metabolic pathways that were most likely to be altered by the observed changes in protein levels, we performed metabolic pathway analysis with Saccharomyces Genome Database (SGD) YeastPathways across all samples from both batches. Several pathways predicted to be most altered were those associated with lipid, amino acid, and nucleotide metabolism (Fig. 5A, Sup. Table 11). We looked more closely at the levels of enzymes with roles in these top pathways and identified key differences between batches. Most enzymes involved in ergosterol biosynthesis were present at higher abundances in Batch 15 than in Batch 1 (Fig. 5B). This was specific to sterol metabolism since enzymes in other lipid metabolism pathways were not significantly different between batches (Sup. Fig. 4D-E). Many of the affected pathways related to sugar and amino acid metabolism contain enzymes Bat1, Bat2, or Pdc5, which are involved in the production of the grainy flavor compound isobutyraldehyde (Fig. 5C). These three enzymes are responsible for the high pathway perturbation scores for pyruvate fermentation, acetoin and butanediol biosynthesis, and degradation of amino acids isoleucine, valine, phenylalanine, tryptophan, and tyrosine: all of these pathways drop out of the top 20 most affected metabolic processes when Bat1, Bat2, and Pdc5 are removed from the dataset, with acetoin and butanediol biosynthesis going from second most perturbed pathway to twenty-second. Interestingly, both Bat2 and Pdc5 abundances were lower in Batch 15, along with a slight decrease in Ald6 which metabolizes isobutyraldehyde to isobutyric acid (Fig. 5D-E).

**Figure 5.**
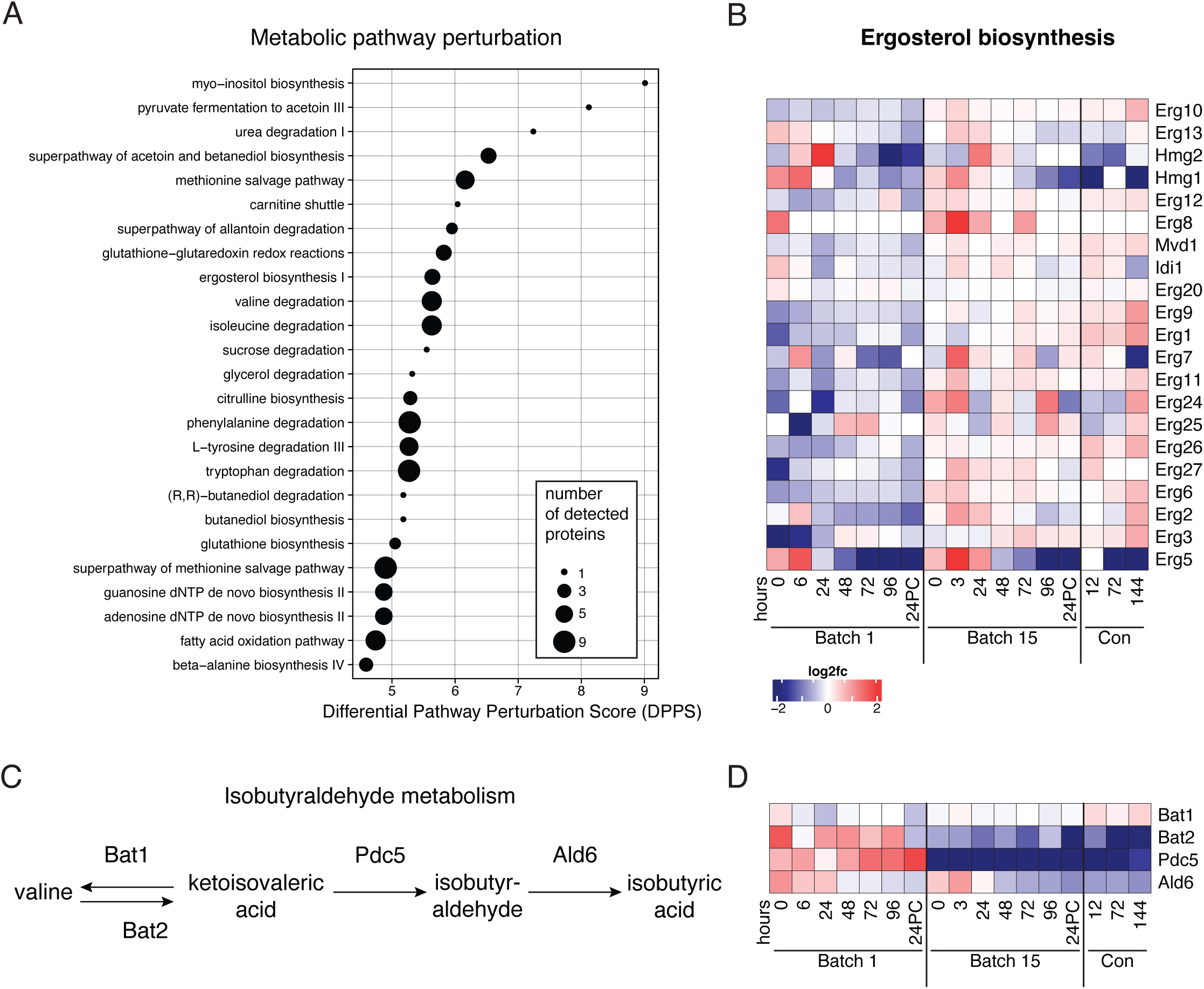
Ergosterol and isobutyraldehyde metabolism differ between early and late batches. A) Top 25 altered metabolic pathways across all samples as determined by SGD Yeast Pathways. B) Changes in ergosterol synthesis enzyme abundances (rows) over both fermentation and conditioning (columns). Color scale represents log_2_ fold change vs. the mean abundance across all time points. C) Steps in the isobutyraldehyde synthesis and metabolism pathway. D) Isobutyrate synthesis is downregulated in Batch 15. Heatmap depicting changes in abundance over time (columns) and enzymes (rows) involved in isobutyrate synthesis.

### Understanding the systems-level regulation of protein modules across brewing

After successful identification of affected metabolic processes by an unbiased method and the high pairwise correlation between protein complexes found in the TCA cycle, we expanded our analysis to globally analyze all annotated protein complexes regulated across fermentation. On intersecting the proteins detected in our dataset with those annotated in the Yeast GFP Fusion Localization database^40^, we detected 74% of all annotated proteins (Sup. Fig. 5A). Our approach broadly sampled proteins across yeast cellular compartments. We detected 75% of the cytoplasmic proteome and nearly all the proteins from the smallest classes such as the lipid particles and those that shuttle between the endoplasmic reticulum and Golgi (Sup. Fig. 5B, Sup. Table 12). We next calculated the pairwise correlations between all proteins in a particular subcellular compartment and plotted distributions of the Pearson correlation coefficient values for every given compartment. Most distributions were centered around zero, indicating that entire compartments were generally not shifting in a concerted manner. Of major cellular compartments, the nucleolar proteome, proteins localized to actin fibers, and proteins that traffic between the ER and Golgi contained well-correlated protein pairs (Sup. Fig. 5C).

Using the Complexome database^41^, we curated a comprehensive set of protein complexes. On average, annotated yeast protein complexes consisted of five members (Sup. Fig. 5D), and we detected members from 79% (492/620) of annotated complexes (Sup. Table 13), on average observing 70% of members within a given protein complex (Sup. Fig. 5E). We generated the pairwise Pearson correlation matrix for all proteins detected across all samples from both batches and identified four well-correlated clusters in the similarity matrix (Fig. 6A). We hypothesized that interacting protein pairs might share similar expression patterns across fermentation. On plotting the distribution of Pearson correlation across all pairs of proteins in our dataset, we indeed found that the distribution of physically interacting protein pairs was more correlated than non-interacting pairs (Fig. 6B). In order to understand the extent of correlation across complexes involved in biological processes, we subset our matrix into individual complexes. The 40S and 60S cytosolic ribosomal subunits were highly correlated compared to their mitochondrial counterparts (Fig. 6C). Chromatin remodeling complexes like RSC, SWI/SNF, and INO80 showed poor pairwise correlations, with their distributions centered around zero (Fig. 6D). Members of RNA polymerase I associated with transcription of rRNAs were highly correlated, unlike mRNA transcribing polymerase II and tRNA transcribing polymerase III (Fig. 6E). However, associated transcriptional co-activators/repressors (TFIID, SAGA, and SLIK) exhibited poor correlation. Membrane transport complexes such as the exocyst and vesicle coat complexes COPI and COPII showed varying degrees of correlation among their members (Fig. 6F). Finally, protein level regulators (26S proteasome) and chaperone complexes (Prefoldin and T-complex) displayed high correlations among their members (Fig. 6G). Therefore, while not all individual protein levels changed over fermentation and repitching, their function could still be impacted by changes in complex partners that show poor correlation.

**Figure 6.**
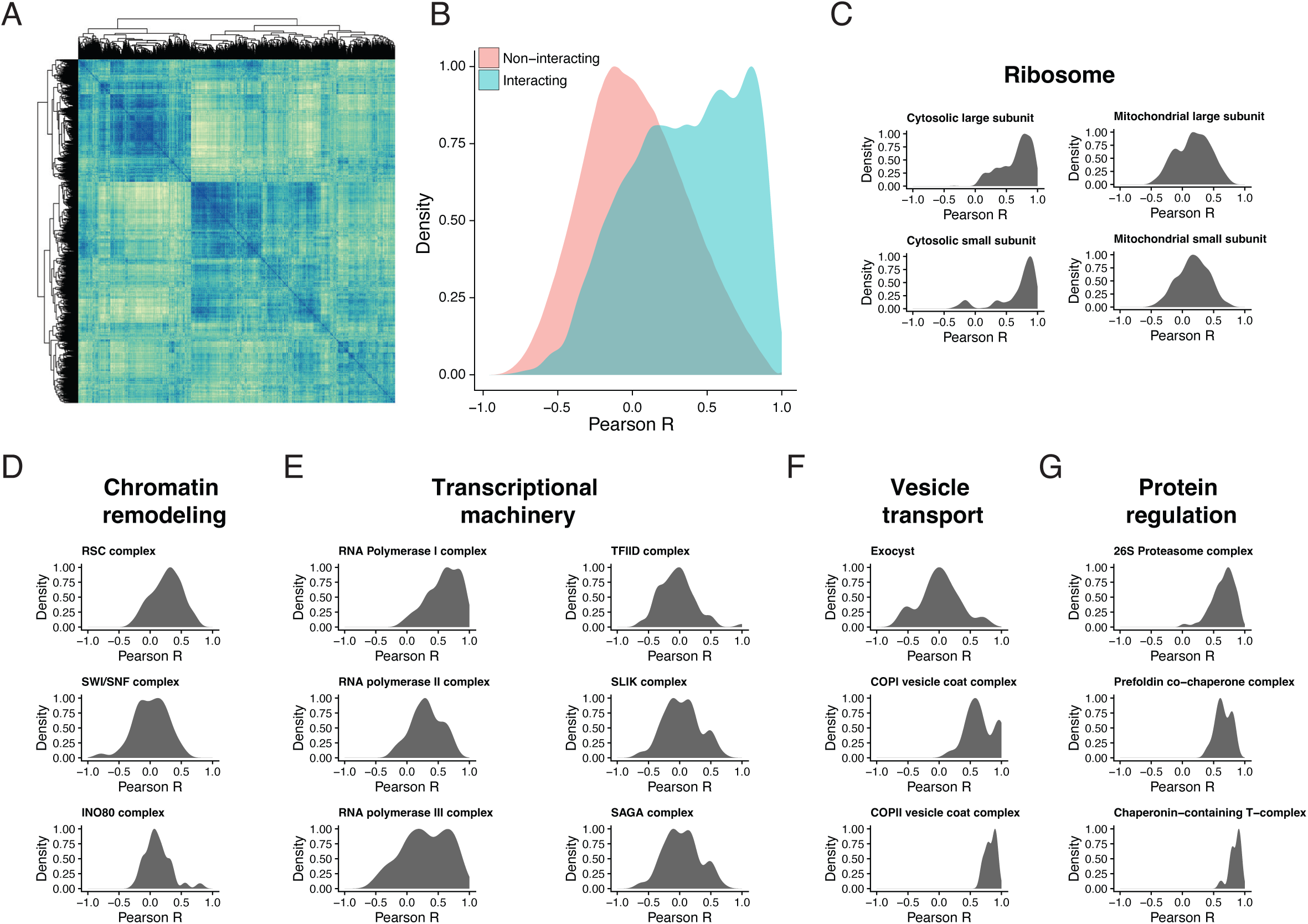
Uncovering protein complex regulation across brewing. A) All by all pairwise protein Pearson correlation matrix where rows and columns represent all 2572 detected proteins across the fermentation dataset. B) Density plot of correlation between known interacting and non-interacting protein pairs (Interaction data curated from EBI complexome^41^). C) Cytosolic ribosomal subunits tend to be more correlated than their mitochondrial counterparts. Density plots of pairwise correlation among protein complex members involved in: D) Chromatin remodeling, E) Transcription, F) Vesicle transport, and G) Protein regulation.

## Discussion

Understanding the molecular changes associated with commercial beer production is crucial to inform the brewing process by guiding strain engineering, identifying molecular characteristics of yeast strains that influence the beer flavor, and optimizing workflows for large scale production. Comprehensively mapping protein dynamics during fermentation offers opportunities to globally identify the enzymes and metabolic pathways responsible for generating the diverse range of flavor compounds in beers and delving into the regulatory mechanisms governing the biochemical processes necessary for fermentation. Furthermore, profiling beer yeast proteomes across successive serial repitching cycles provides a unique lens into the evolutionary and physiological dynamics and adaptive responses of yeast populations during brewing.

Here, we characterized the genome of a Hefeweizen ale brewing strain, Wyeast 3068, and the changes in its proteome throughout fermentation in a commercial brewing setting. We characterized a fermentation time course across two batches separated by 14 repitches to elucidate the impact of serial repitching on the brewing yeast proteome. While previous work has characterized the genetic changes in brewing yeast genomes^19^, traced ecological origins of beer brewing strains^10–12^, and mapped the residual proteins in beer^22–24, 38, 42^, our work systematically profiled the proteomic changes in an ale yeast strain across entire sets of commercial fermentation cycles using shotgun mass spectrometry. From this study, we have created a comprehensive dataset cataloging 2572 yeast proteins across 17 timepoints during industrial beer brewing. Our data reveal global trends in protein expression changes over brewing cycles as well as across serial repitching, a practice widely adopted by many breweries but understudied with respect to molecular changes in the fermenting yeast populations.

Our genetic characterization of Wyeast 3068 showed deletions of many genes as expected for ale yeast: flocculation genes vary between brewing strains^43^ and can be lost in aged brewing yeast^44^, less efficient carbohydrate transporters are lost in some brewing strains^45^, *ASP3* has been lost in many *S. cerevisiae* isolates^46^, and transposition events and copy number variations of Ty elements are common in industrial yeast strains^47^. Previous studies have demonstrated that genome evolution in brewing yeast strains does occur across the repitching process as selected populations have adapted to the fermentation environment^19^. While it is beyond the scope of this work, the mechanism of strain adaptations warrant further studies such as matched genomic data across fermentation cycles and serial repitching. It remains to be seen to what degree the changes we observed in protein abundances arose from genetic mechanisms (i.e. deletions or duplications) versus gene expression regulation at the RNA and post-translational levels; protein activities might also be changing independently of abundances, such as by post-translational modification or allosteric or feedback regulation, none of which we have attempted to measure here.

By focusing on proteomic changes over the course of fermentation, we found that the largest number of proteins whose abundances significantly change occur within the first 24 hours of fermentation. Sampling more time points in the first day of fermentation would provide finer resolution of the protein expression changes. Importantly, whether these changes are purely associated with growth versus immediate acclimatization to the fermentation tank merits further investigation. Strikingly, protein translation machinery and ribosome biogenesis peaked three to six hours after the start of fermentation before being downregulated. These trends suggest that the yeast strains are primed for protein synthesis before entering strictly anaerobic conditions later in fermentation. It is interesting to note that while certain protein clusters specific to the conditioning tank do not significantly represent a GO biological process, they tended to consist of mitochondrial ribosomal subunits and those associated with protein translation. Additionally, they also consisted of a handful of uncharacterized proteins. Further studies characterizing these sets of proteins will shed light on the molecular processes involved in the conditioning process.

Broadly, we found protein trends across fermentation to be similar before and after serial repitching barring some marked differences. For example, ergosterol synthesis enzymes were elevated in Batch 15. Ergosterol can help counter the stress of hypoxic environments, enabling normal growth and flocculation necessary for brewing^48^. This may be caused by the selection for yeast that are better able to survive hypoxic conditions due to increased ergosterol synthesis, suggesting ergosterol biosynthesis as a potential target for beer yeast strain engineering. Alternatively, it may be indicative of cellular stress programs that increase ergosterol production, which could be mitigated by ergosterol supplementation to reduce stress and confer better growth behavior for late-pitch yeast^49^. The other striking difference between batches was in isobutyraldehyde synthesis enzymes Bat2 and Pdc5, which were less abundant in Batch 15 than in Batch 1. Isobutyraldehyde is associated with a grainy flavor profile, which is desirable in some beers and considered an off-flavor in others. In winemaking, deletions of *BAT2* and *PDC5* can be used to reduce the production of undesirable fusel alcohols^50^. The observed changes in Bat2 and Pdc5 abundances between early and late batches could contribute to inconsistencies in flavor over many repitches. Why these enzymes change over batches remains to be explored.

While we measured the abundances of thousands of proteins across fermentation cycles, our approach is limited to only measuring protein levels and not post-translational modifications or metabolite changes across fermentation. Many metabolic enzymes that regulate important processes in fermentation are not only regulated by absolute levels but also by modifications such as phosphorylation and glycosylation. Many proteins for which we did not observe changes in absolute levels could still have altered activity over fermentation due to post-translational modifications. For example, phosphofructokinase and pyruvate decarboxylase are regulated by phosphorylation^51, 52^. Additionally, it is known that secreted yeast glycoproteins contribute to the proteome of beer, but how they change over time has not been investigated^53^. A recent study identified changes in the phosphoproteome of yeast during diauxic shift^34^, so further studies such as phosphoproteomics over fermentations and before and after repitching would likely identify other important pathways regulated in brewing. Coupling measurement of yeast enzyme levels to metabolite levels in the fermentation tank could give a comprehensive view of how yeast biology is altered in brewing and how that impacts the fermentation product.

Our study provides a systems biology view of the molecular processes that underlie beer brewing. By analyzing how changes in protein levels alter protein complexes and biochemical pathways, we observed that interacting protein pairs are correlated across samples, suggesting that many yeast cellular modules are co-regulated during brewing. Future studies characterizing the protein-protein interactions using proximity labeling^54, 55^ or fractionation^56^ with mass spectrometry across brewing will shed light on the dynamic regulation of protein complexes across fermentation.

Finally, we believe that this resource serves as a comprehensive catalog of fermentation-based protein changes, and we have made it available for exploration on an interactive web interface (https://brewing-yeast-proteomics.ccbb.utexas.edu/). Further analysis of our data and future studies of proteins, post-translational modifications, and metabolite changes across fermentation and repitching will aid beer yeast strain engineering, optimization of brewing workflows, and study of trends that underlie domestication processes.

## Methods

### Strain

Wyeast 3068, Weihenstephan Wheat yeast.

### Hefeweizen brewing

Commercial fermentation with Wyeast 3068 was conducted at Live Oak Brewing Company, Austin, Texas, USA. For Batch 1, 32 liters of yeast cultured on rich media supplied by Wyeast was inoculated into 100 gal of wort; 24 hours later this was inoculated into 60 bbl of wort. Fermentation proceeded at 20°C in a horizontal tank for four to five days, after which it was cold crashed via glycol jacket heat exchange to 4.4°C to sediment the yeast. The beer was then separated from the yeast and moved to a conditioning tank to be held at cold temperatures for two weeks. Directly after this transfer, the yeast from the fermenter is harvested to be used in the subsequent batch, pitched either that day or the following day. Fermentations for batches 2-14 (between batches 1 and 15) occurred over the same four to five day duration, but now fermenting 180 bbl with 65 liters of yeast. All batches were used to produce Hefeweizen ale and both batches sampled for proteomics were subjected to the same wort and fermentation conditions.

### Whole genome sequencing

For genomic DNA isolation, a glycerol stock of Wyeast 3068 strain received from the brewers was streaked onto rich media (YPD) agar plates and incubated for 3-5 days. A single colony was isolated and cultured for DNA isolation. Genomic DNA extraction was performed using Zymo YeaStar Genomic DNA Isolation kit as per manufacturer recommendations. Genomic DNA was sheared to an average length of 410 bp before sequencing on an Illumina NextSeq 500 platform. Sequences were first analyzed using FastQC to assess overall quality. The 1.6 million read pairs were then mapped to the *Saccharomyces cerevisiae* S288C reference genome R64 (sacCer3) using bowtie2 (v 2.2.6) in local alignment mode. We observed a high overall mapping rate of 92% with low rates of duplication (0.5%) and reads mapping to multiple locations (4.2%). A moderate rate of indel detection (13.4%) and the relatively low 68.8% of reads mapped as proper pairs suggested this commercial yeast differs from the lab strain in some structural ways. Sequencing reads are deposited in the NCBI Sequence Read Archive (SRA) under BioProject PRJNA1011390.

### Copy number analysis

Copy number variation was calculated by read coverage over 1000 base pair sliding windows, as described in Large et al^19^. Single-gene deletions were identified using CNVnator^57^ and confirmed by viewing bam files in IGV^58^. The list of affected genes was input into the SGD Gene Ontology Term Finder version 0.86 (https://www.yeastgenome.org/goTermFinder) to identify process ontology aspects.

### Loss of heterozygosity (LOH) analysis

To detect genomic regions of the brewing yeast that experienced the loss of heterozygosity, we performed variant calling and allele frequency analyses using the CLC Genomics Workbench 11.0 NGS toolkit platform (Qiagen). After importing paired-end Illumina FASTQ read files, Nextera adapters were trimmed, and the trimmed reads were then mapped to the *S. cerevisiae* R64 (sacCer3) reference genome, with no masking. Mapping was random, using the following parameters: mismatch cost 2, insertion and deletion cost 3, length fraction 0.5, and similarity fraction 0.8. We then used Workbench’s Basic Variant Detection 2.0 program to identify variants and to calculate the frequency of each variant allele, using the following parameters: ignore broken pairs; ploidy 4; minimum coverage 10; minimum count 2; minimum frequency 0%. Variants were spot-checked on the pile-up viewer of the mapped reads, with all confirmed to be correctly called. After removing the called variants for the mitochondrial genome, the resulting output file was ordered by chromosome number, and whole-genome (concatenated) positions were assigned for each variant. The allele frequencies of the variants were then plotted against the whole-genome position (Sup. Fig. 1B).

### Sample preparation for proteomics

Yeast proteins for mass spectrometry were isolated using a previously described protocol^59^. Briefly, samples were obtained directly from beer fermentation tanks at Live Oak Brewing Company in Austin, Texas, USA. For every time point, two replicates were independently sampled from the fermentation tank by collecting 1 liter per replicate. Cell pellets were harvested by centrifuging beer at 8000g for 5 min followed by 2-3 washes in ice cold PBS. Cell pellets were resuspended in Digestion Buffer (50mM Tris, 2mM CaCl_2_) and lysed by bead beating with glass beads for 1 minute cycles repeated three times. The whole-cell lysate was separated from the beads and then mixed 1:1 with 2,2,2-trifluoroethanol (TFE). Samples were then reduced by incubation with 5mM tris(2-carboxyethyl)phosphine (TCEP solution, Pierce) at 60°C for 40 minutes. Reduced samples were alkylated by incubation with 15mM iodoacetamide at room temperature for 30 minutes. Excess iodoacetamide was quenched by the addition of 7.5mM dithiothreitol (DTT). Following quenching, samples were diluted 10-fold using digestion buffer and subjected to proteolytic digestion with 2µg trypsin for 5 hours at 37°C. Tryptic digestion was quenched with 1% formic acid, and samples were concentrated using vacuum centrifugation to reduce the total sample volume to less than 300µl. Digested samples were cleaned using HyperSep C18 SpinTips (Thermo) according to the manufacturer’s protocol. Eluted peptides were briefly dried by vacuum centrifugation, then resuspended in 5% acetonitrile and 0.1% formic acid.

### LC-MS/MS analysis

Tryptic peptides were separated by reverse phase chromatography on a Dionex Ultimate 3000 RSLCnano UHPLC system (Thermo Scientific) with an Acclaim C18 PepMap RSLC column using a 3-42% acetonitrile gradient over 60 minutes. Peptides were eluted directly into a Thermo Orbitrap Lumos mass spectrometer by nano-electrospray. Data-dependent acquisition (DDA) was applied, with precursor ion scans (MS1) collected by FTMS at 120,000 resolution and HCD fragmentation scans (MS2) collected in parallel by ITMS with 3-second cycle times. Monoisotopic precursor selection and charge-state screening were enabled, with ions > +1 charge selected. Dynamic exclusion was applied to selected ions +/-10 ppm for 30 seconds. Raw mass spectrometry data have been deposited on MassIVE (MSV000092793).

### Proteome database searching and analyses

Raw mass spectrometry data was processed using Proteome Discoverer 2.2, MaxQuant, or converted to mascot generic files (.mgf) using MSConvert to be analyzed by SearchGui and PeptideShaker. Mass spectra were searched against a protein sequence database containing reversed decoy sequences comprising the *S. cerevisiae* reference proteome (UniProt OX: 559292) and a list of common protein contaminants (MaxQuant). All searches were restricted to fully tryptic peptides only, allowing up to two missed cleavages. A precursor tolerance of 5 ppm and fragment mass tolerance of 0.5 Da were used. Static modifications of carbamidomethyl cysteine and dynamic modifications of oxidized methionine and protein N-terminal acetylation and/or methionine-loss were considered. High-confidence peptide spectrum matches (PSMs), peptides, and proteins were all filtered at a false discovery rate of <1%.

Protein abundances were calculated using three different metrics: (i) estimating parts per million abundance from PSM counts^29^ (ii) Intensity-based Absolute Quantification (iBAQ) and (iii) Label-Free Quantitation (LFQ)^30–32^. For ppm-based protein abundance, mascot generic files were then searched against MS-GF+, OMSSA, and X!Tandem databases with default settings for each database using SearchGUI (version 3.2.20)^60^ and data were analyzed using PeptideShaker version 1.16.12^61^. PeptideShaker report files were parsed to generate a matrix of unique validated PSMs for each protein across the fermentation time course. To normalize the PSM counts, we converted the count matrix for a given protein into parts per million (ppm) using an approach previously described^29, 62^. Briefly, unique peptides were trypsin digested *in silico* and filtered to only those 7-40 amino acids in length. Next, a correction factor was calculated from the sum of the total length of peptides in this range per protein. Detected peptide PSMs were multiplied by the peptide length, and correction factor, multiplied by 1,000,000, and divided by the experiment total to get parts per million. iBAQ and LFQ-based protein abundances were calculated using MaxQuant^32^ based on extracted ion-chromatography (XIC) feature intensities.

To compute similarity matrices across time points, pairwise Pearson correlation was calculated between each pair of sampled time points. Since the LFQ values had the highest correlation between replicates and are corrected for technical variation between samples, we proceeded with LFQ abundances for all subsequent analyses. LFQ intensities for a protein were across technical replicates to maximize the number of proteins detected across all samples.

### Clustering, differential expression, and GO term analysis

Raw LFQ intensity matrices from the MaxQuant output were filtered to ensure that each protein was detected with at least two unique peptides. Since LFQ intensity values are log-normally distributed, we log_2_-transformed the data to obtain a normal distribution of intensities and median centered to ensure equal variances across samples. Only proteins that were detected in both replicates of a particular timepoint were considered for downstream analyses. Pairwise significance testing across time points was performed using a standard t-test with p-values adjusted using the Benjamini-Hochberg (BH) procedure with a false discovery rate of 5%. Enriched proteins were further filtered to a log_2_ fold change of 1 or greater. The above procedures were carried out on the online version of ProVision^63^.

For hierarchical clustering, log_2_ protein abundances were normalized to row mean (Sup. Table 6) and proteins that did not change more than two-fold over the mean in any time point were filtered out. Hierarchical row clustering was performed by Morpheus (https://software.broadinstitute.org/morpheus) using an average linkage method. We identified eight clusters at a node depth of three that contained 36-748 proteins. Clusters with over 500 proteins were further subset to a node depth of five, and resultant subclusters contained 59-387 proteins (Sup. Table 7). GO enrichment analysis was performed using clusterProfiler^35^ for each cluster and sub-cluster to determine significantly enriched proteins by comparing protein changes in a pairwise manner across all time points. GO term enrichments were filtered to a multiple hypothesis testing FDR of 5% and BH adjusted p-value of less than 0.05 (Sup. Table. 8).

### Metabolic pathway analysis

Prediction of metabolic pathways affected by changes in protein levels over time across both batches was performed using the cellular overview tool on SGD Yeast Pathways. Log_2_ protein abundances normalized to row mean (Sup. Table 6) with gene name were uploaded onto the Omics Viewer, and output set to a table of 100 top-scoring pathways (Sup. Table 11). Differential Pathway Perturbation Scores (DPPS) were calculated as described in the tool. Briefly, a differential reaction perturbation score (DRPS) is calculated from the maximum of differences between samples for all entities associated with a given reaction. For a pathway, the square of each DRPS is summed and divided by the total number of reactions in the pathway. The DPPS is the square root of this value, representing the maximal perturbation of the pathway between samples.

### Pathway, protein complex, and subcellular location analysis

Yeast pathways were curated from SGD^5, 64^. Yeast protein complexes were obtained from the EBI complex portal^41^. For every protein complex, the fraction of members detected was computed (Sup. Table 13). Since the EBI complex portal manually curates protein complexes across the multiple protein databases, there existed multiple versions or variants of a particular complex. In these cases, for our analyses, we considered these to be discrete complexes. We calculated Pearson correlation across all the fermentation time points for all possible protein pairs in the dataset and subset them based on whether they were interacting or not. Protein subcellular localization data were curated from the yeast GFP fusion localization database^40^. In cases where a protein localized to more than one location, these proteins were annotated in both subcellular locations or organelles to prevent loss of information (Sup. Table 12).

## Acknowledgments

The authors would like to thank the University of Texas at Austin Genome Sequencing and Analysis Facility (GSAF) for assistance with NGS library preparation and sequencing. The authors would like to thank Kevin Drew (University of Illinois, Chicago), Devin Schweppe (University of Washington), and members of the Marcotte and Dunham laboratories for insightful discussions and helpful feedback.

## Funding Sources

This material is based in part upon work supported by the National Science Foundation under Cooperative Agreement No. DBI-0939454. Any opinions, findings, conclusions or recommendations expressed in this material are those of the author(s) and do not necessarily reflect the views of the National Science Foundation. R.K.G. acknowledges support from the University of Washington’s Center for Multiplex Assessment of Phenotype postdoctoral fellowship. J.O.A. acknowledges support from University of Washington Biological Mechanisms of Healthy Aging training grant. R.C.G. was supported by the National Institute of General Medical Sciences of the National Institutes of Health under award F32 GM143852 and by the Momental Foundation. M.L. was supported by the Swiss National Science Foundation under grant P5R5PB_211122. E.M.M. acknowledges support from the National Institute of General Medical Sciences (R35 GM122480), Army Research Office (W911NF-12-1-0390), and Welch Foundation (F-1515). The authors acknowledge the Texas Advanced Computing Center (TACC) at The University of Texas at Austin for providing high-performance computing resources that have contributed to the research results reported within this paper.

## Competing Interests

E.M.M. is a co-founder, shareholder, and scientific advisory member of Erisyon, Inc., which played no role in this work. D.K. is an employee, T.P. is a former employee, and H.M.E is a founding member of Live Oak Brewing Company. The other authors declare no competing interests.

## Author Contributions

R.K.G. (conceptualization, methodology, data curation, formal analysis, investigation, visualization, writing-original draft), R.C.G. (formal analysis, writing-original draft, visualization), J.O.A. (formal analysis, writing-original draft, visualization), B.D. (formal analysis, writing-original draft), D.R.B. (conceptualization, methodology, analysis, investigation, visualization), A.B. (investigation, formal analysis), M.L. (software), V.D. (software), P.J. (formal analysis), D.K. (resources), T.P. (resources), H.M.E. (resources), E.M.M. (funding acquisition, conceptualization, project administration, supervision, resources, writing-review and editing), M.J.D. (funding acquisition, project administration, supervision, writing-review and editing).

---

**Supplementary Figure 1.**
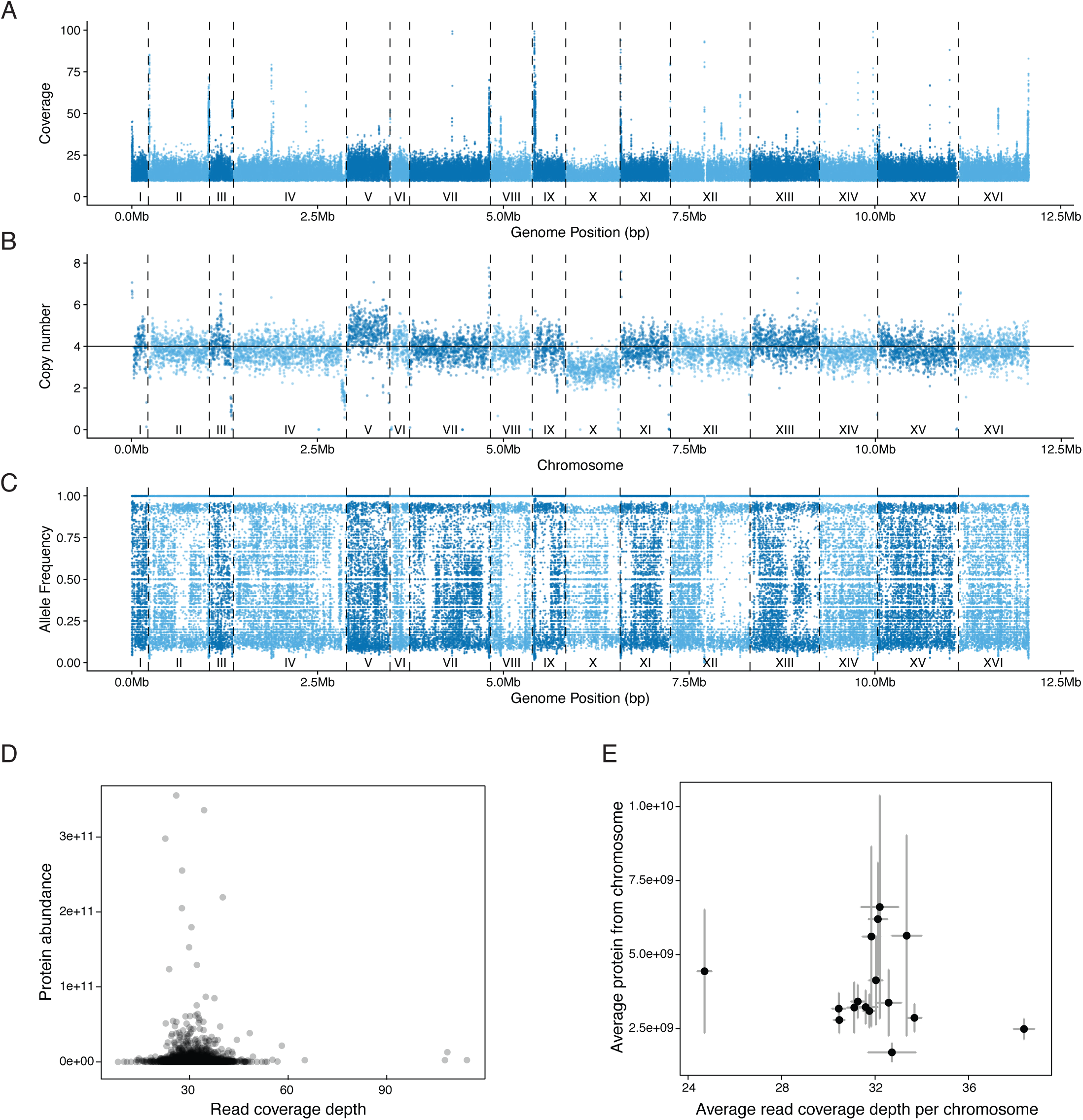
Genomic characteristics of Wyeast 3068. A) Mean sequencing coverage across the Wyeast 3068 genome. Each dot represents the mean coverage (y-axis) of a particular locus along the genome position (x-axis). B) Local copy number as an average read depth over 1000bp windows normalized to the known ploidy of 4n. C) Allele frequencies (y-axis) of called variants. (Note that genome positions in A, B, and C have been concatenated for whole-genome display). D) Comparison between read depth (x-axis) at each gene and level of the corresponding protein abundance (y-axis). Adjusted R^2^=0.00008246, p≤0.2703. E) Comparison between average read depth/chromosome (x-axis) and average LFQ abundance of corresponding protein by chromosome (y-axis), adjusted R^2^=-0.04194, p≤0.5392.

**Supplementary Figure 2.**
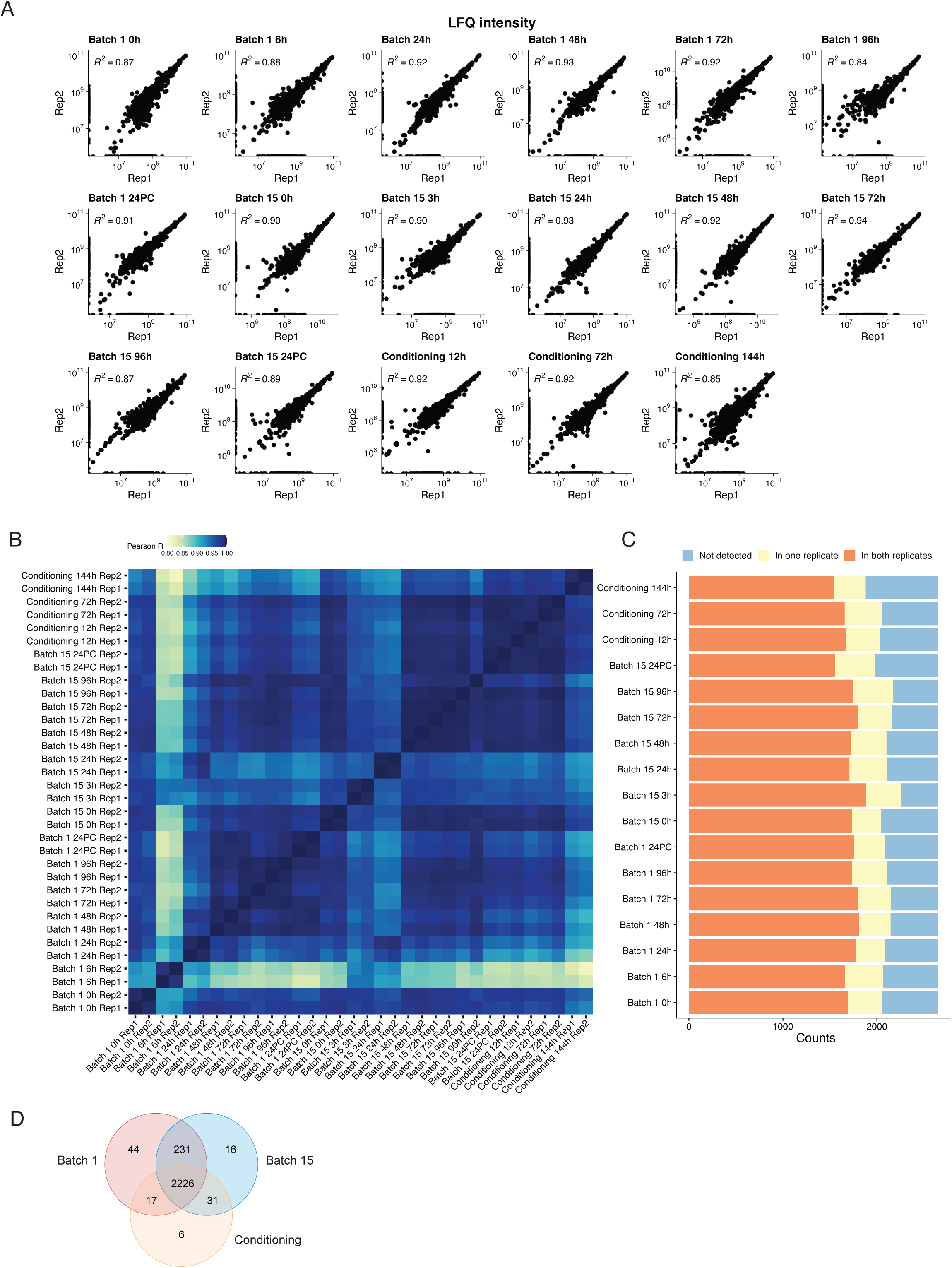
Fermentation proteomics summary statistics. A) Scatterplots depicting the correlation between replicates 1 and 2 for each fermentation time point assayed. Axes indicate the log-transformed observed LFQ intensity calculated using MaxQuant. B) Pairwise similarity matrix showing the extent of correlation across all sampled replicates and time points in the dataset. C) Barplots summarizing the number of proteins detected in each time point colored by detection in both (orange), one (yellow), or undetected (blue). D) Venn diagrams summarizing the overlap of proteins detected across Batch 1 (red), Batch 15 (blue), and the Conditioning tank (orange).

**Supplementary Figure 3.**
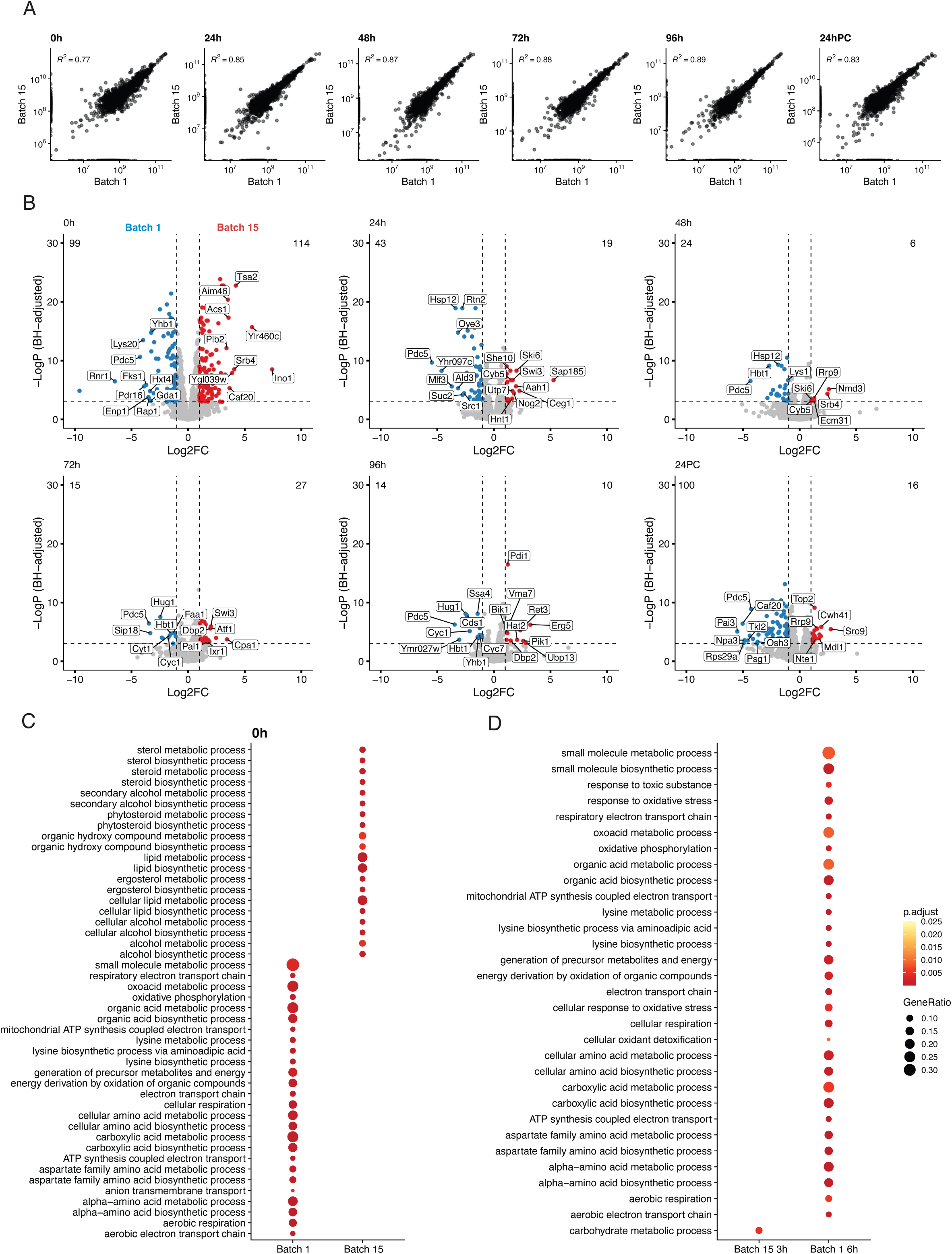
Comparing differentially expressed proteins across shared time points between batches 1 and 15. A) Scatter plots of summed LFQ intensity across replicates comparing time points sampled in both Batch 1 (x-axis) and Batch 15 (y-axis) brewing cycles. B) Differentially expressed proteins across matched time points (0h, 24h, 48h, 72h, 96h, and 24PC) between Batches 1 and 15 depicted using volcano plots with log_2_ fold change (x-axis) and Benjamini-Hochberg adjusted p-value (y-axis). C) Dotplots with the top enriched GO terms for the matched starting time point across Batch 1 and Batch 15. D) Dotplots with top enriched GO terms across Batch 15 3h and Batch 1 6h time points. GO term analysis was performed on the biological function terms with a 5% FDR threshold and filtering terms to an adjusted p-value (Benjamini-Hochberg correction) of <0.05. Sizes of dots correspond to the ratio genes detected to the total genes annotated for a particular GO term.

**Supplementary Figure 4.**
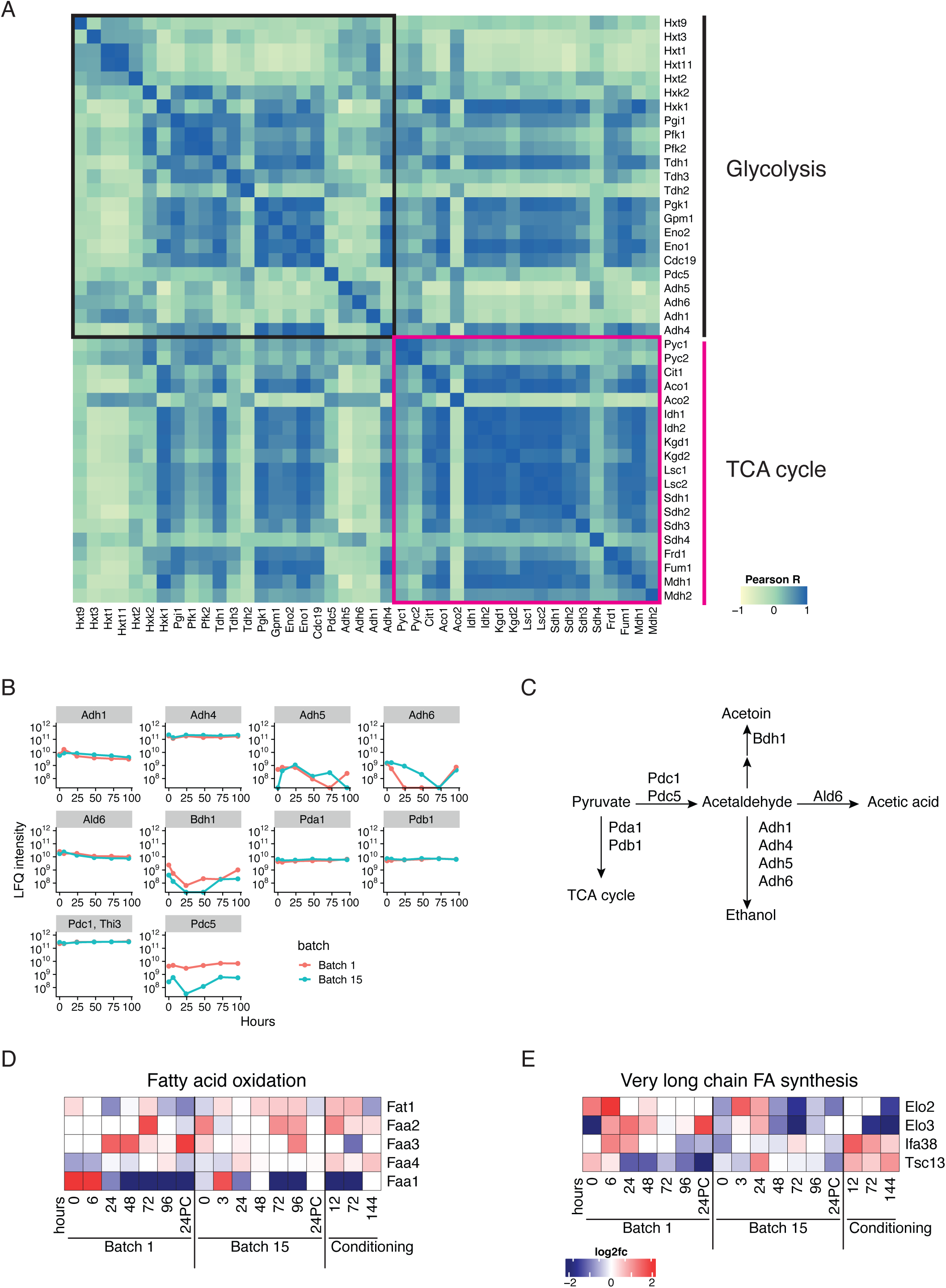
Cataloging abundance changes in metabolic pathways. A) Pairwise Pearson correlation between proteins involved in the glycolysis and tricarboxylic acid (TCA) cycle pathways. B) LFQ protein abundance (y-axis) as a function of time (x-axis) for enzymes involved in pyruvate metabolism. Time points colored by batches 1 (red) and 15 (blue) C) Steps in pyruvate metabolism in yeast. D) Changes in fatty acid oxidation and E) very long chain fatty acid synthesis enzyme abundances, over both batches and during final conditioning, as log_2_ of row mean normalized abundance.

**Supplementary Figure 5.**
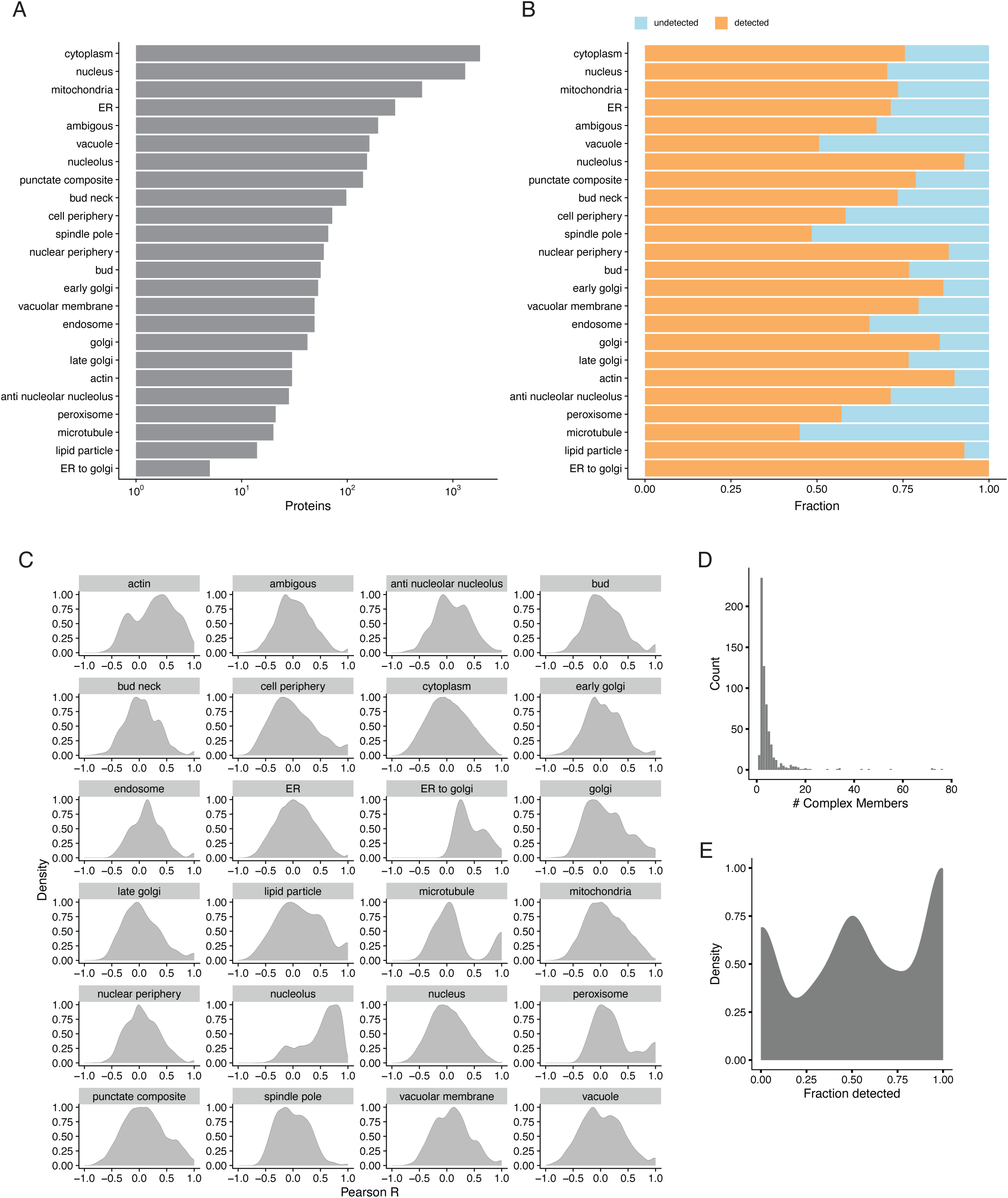
Subcellular proteome analysis. A) Barplots representing the numbers of yeast proteins (x-axis) annotated by subcellular/organellar location (y-axis). Data curated from the Yeast GFP Fusion Localization database^40^ and B) Fraction of proteins in each subcellular/organellar location detected in at least one time point across the brewing time course. C) Densities of all Pearson correlation values calculated across all pairs of proteins across each annotated subcellular location and organelles. D) Histogram of the distribution of yeast protein complex sizes. Data obtained from Yeast complexome. E) Density plot showing the fraction of yeast protein complexes detected in the fermentation time course. Protein complex data curated from the EBI Complexome database^41^.

## Notes

### Competing Interest Statement

The authors have declared no competing interest.

### Summary of Updates

This manuscript has been updated to fix typos in and formatting in the figures

